# Methionyl-tRNA synthetase inhibitor has potent *in vivo* activity in a novel *Giardia lamblia* luciferase murine infection model

**DOI:** 10.1101/822924

**Authors:** Samantha A. Michaels, Han-Wei Shih, Bailin Zhang, Edelmar D. Navaluna, Zhongsheng Zhang, Ranae M. Ranade, J. Robert Gillespie, Ethan A. Merritt, Erkang Fan, Frederick S. Buckner, Alexander R. Paredez, Kayode K. Ojo

## Abstract

**Objectives:** Methionyl-tRNA synthetase (MetRS) inhibitors are under investigation for the treatment of intestinal infections caused by *Giardia lamblia*. To properly analyze the therapeutic potential of the MetRS inhibitor **1717**, experimental tools including a robust cell-based assay and a murine model of infection were developed based on novel strains of *G. lamblia* that employ luciferase reporter systems to quantify viable parasites.

**Methods:** Systematic screening of Giardia-specific promoters and luciferase variants led to the development of a strain expressing the click beetle green luciferase. Further modifying this strain to express NanoLuc created a dual reporter strain capable of quantifying parasites in both the trophozoite and cyst stages. These strains were used to develop a high throughput cell assay and a mouse infection model. A library of MetRS inhibitors was screened in the cell assay and **1717** was tested for efficacy in the mouse infection model.

**Results:** Cell viability in *in vitro* compound screens was quantified via bioluminescence readouts while infection loads in mice were monitored with noninvasive whole-animal imaging and fecal analysis. Compound **1717** was effective in clearing mice of *Giardia* infection in 3 days at varying doses, which is supported by data from enzymatic and phenotypic cell assays.

**Conclusions:** The new *in vitro* and *in vivo* assays based on luciferase expression by engineered *G. lamblia* strains are useful for the discovery and development of new therapeutics for giardiasis. MetRS inhibitors, as validated by **1717**, have promising anti-giardiasis properties that merit further study as alternative therapeutics.

## Introduction

*Giardia lamblia* infections are among the most common causes of chronic diarrhea in children in resource-limited environments. New therapeutics are needed to address issues with existing therapies including resistance, toxicity, and reduced efficacy. No vaccines have been developed for clinical use, so case management depends solely on antimicrobial chemotherapy.^1^ Current therapies approved by the Food and Drug Administration for giardiasis include metronidazole and tinidazole.^2–4^ However, approximately 20% of clinical cases involve metronidazole- (and presumably tinidazole-) resistant *Giardia.*^5^ Second-line drugs such as albendazole, nitazoxanide, furazolidone, and paromomycin generally have lower efficacy rates and/or potentially dangerous side effects.^6^ All of these factors necessitate the development of a new therapeutic, which requires various experimental tools for screening and verification of efficacy of potential leads.

While experimental methods suitable for drug screening against *G. lamblia* have been described in literature,^7^ *in vivo* infection models in adult mice can benefit from an improved method of monitoring infection. This deficiency of the *in vivo* model can be explained in part by earlier studies showing instability in the expression of transgenes in *Giardia* compared to many other eukaryotic cells.^8^ The instability leads to variable expression as *Giardia*, by an unknown mechanism, downregulates transgenic markers.^8^ We recently described a luciferase-based reporter system in *G. lamblia* WBC6 trophozoites for quantitative fluorescence readouts from a red-shifted firefly luciferase reporter gene (PpyRE9h) under the control of the β-tubulin promoter (pβTub).^9, 10^ While the bioluminescent output of *G. lamblia* WBC6:pβTub∷PpyRE9h was sufficiently stable for 24-hour *in vitro* assays, it lacked long-term stability over multiple passages and proved unsuitable for the mouse model of infection. In this study, we describe the development of a new strain of *G. lamblia* using a click beetle green luciferase (CBG99) under a short glutamate dehydrogenase promoter (pGDHS) that can be used for *in vitro* screens of compound libraries and *in vivo* efficacy assays. A second strain was engineered by endogenously tagging one copy of the cyst wall protein 1 (CWP1) gene with NanoLuc (Nluc) in the *G. lamblia* WBC6:pGDHS∷CBG99 strain, establishing a second developmentally induced reporter that specifically measures cyst quantity. The strains can be visualized and measured in an animal model of infection using noninvasive imaging to evaluate experimental drug effects in *Giardia*-infected animals. The developed mouse model was validated using the standard metronidazole treatment and subsequently was used to confirm that *G. lamblia* methionyl-tRNA synthetase (MetRS) enzyme inhibitor **1717** is a potential therapeutic alternative for treatment of clinical giardiasis.

## Materials and Methods

### Plasmid construction and G. lamblia transfection

To optimize luminescence of *Giardia* for *in vitro* and *in vivo* experiments, various combinations of luciferase genes with *Giardia-*specific promoters were tested. Twelve gene constructs were generated: three containing PpyRE9h driven by three *Giardia* promoters, five containing various luciferase genes driven by the long glutamate dehydrogenase promoter (pGDHL), and four containing CBG99 driven by four *Giardia* promoters. The three PpyRE9h constructs were driven by pGDHL, the short glutamate dehydrogenase promoter (pGDHS),^11^ and the ornithine carboxytranferase promoter (pOCT).^12^ These were amplified by PCR using the primers found in Table 1 and digested with EcoRI and XbaI. The promoter fragments were cloned into the EcoRI-XbaI site of integration vector pPACV-integ.^10, 13^ For the luciferase systems driven by pGDHL, the coding regions of the various luciferase genes were amplified with the primers from Table 1 and digested with XbaI and PacI. The XbaI-PacI fragments were cloned downstream of the promoter in the pPACV-integ vector’s XbaI-PacI cloning site. Selected luciferase genes included standard firefly luciferase (Fluc), firefly luciferase 2 (Luc2), red click beetle luciferase (CBR), enhanced green-emitting luciferase (Eluc), and CBG99. To generate the CBG99 constructs, the pGDHL, pGDHS, pOCT, and pβTub were amplified with primers from Table 1, digested using EcoRI and XbaI, and cloned into the EcoRI and XbaI digested vector pPACV-integ. To generate the CWP1-NLuc construct, the CWP1 (GL50803_5638) and NLuc genes were amplified with the primers from Table 1 and digested with XbaI-BamHI and BamHI-EcoRI, respectively, then cloned into the XbaII and EcoRI digested pKS-NEO vector.

**Table 1.**
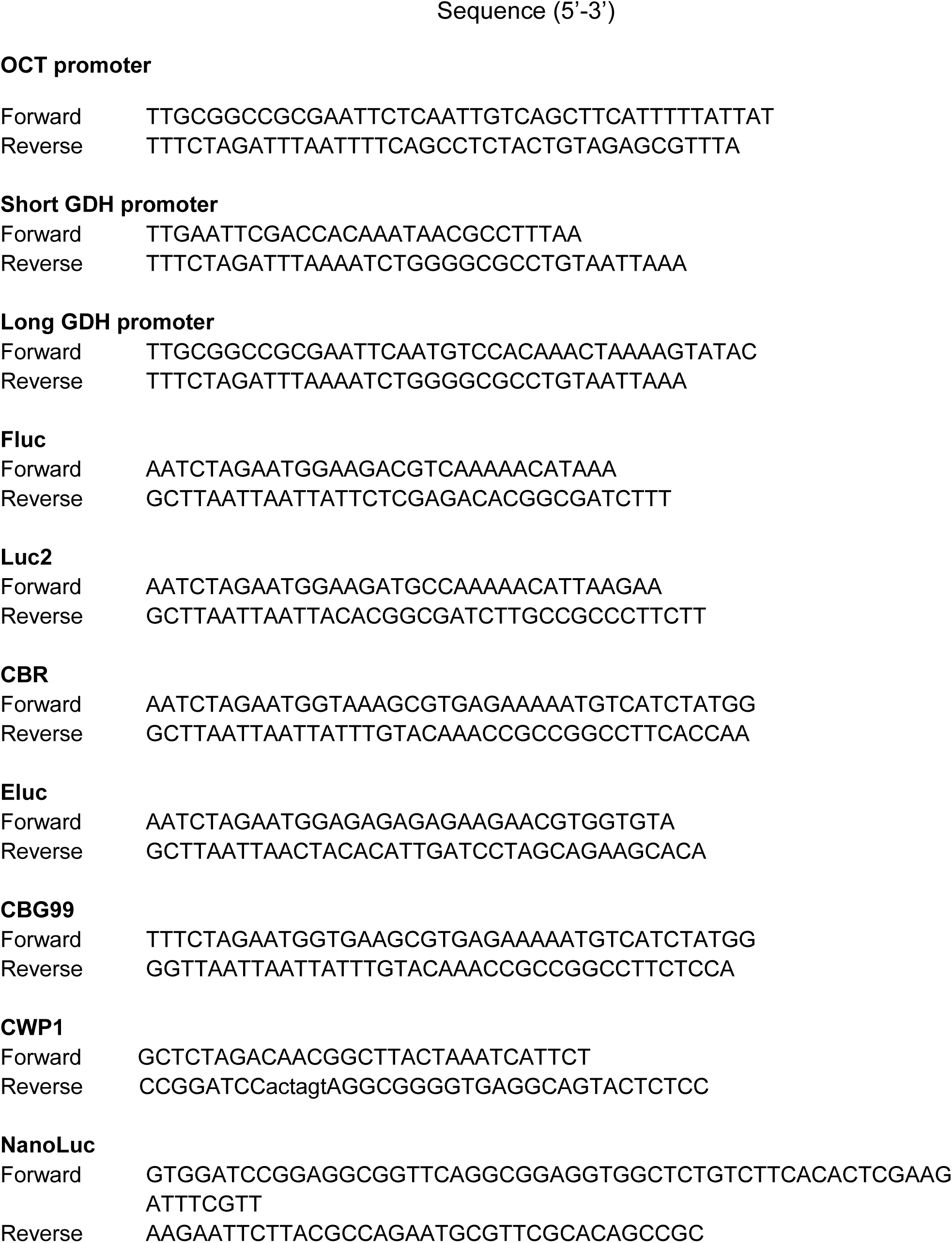
List of primers used in this study.

For integration, 30 µg of the pPACV-integ constructs was linearized with SwaI overnight at 25°C while CWP1-NLuc/pKS-NEO construct was linearized with StuI overnight at 37°C. They were then precipitated with ethanol and incubated with 300 µL of chilled *G. lamblia* cells (~13×10^6^ cells/mL) for 30 minutes before electroporation (Bio-Rad GenePulser X at 375 V, 1000 µF, 750 ohms). After electroporation, cells were incubated on ice for 10 minutes, then transferred to fresh media at 37°C. Transfectants were selected with puromycin after overnight recovery.

### Parasite cultures

*G. lamblia* WBC6:pβTub∷PpyRE9h and wild type (WBC6, ATCC 50803) trophozoites were the starting strains used in these studies.^10^ These and all the newly transfected strains were grown in TYI-S-33 medium supplemented with 10% bovine serum and 0.05 mg/mL bovine bile without (wild type WBC6) or with (transfected strains) antibiotic selection pressure of 32 µg/mL puromycin dihydrochloride (Gibco, Dublin, Ireland).^10, 14^ Cultures were incubated at 37°C in 16 mL Falcon round bottom polystyrene test tubes (Corning Inc., Corning, NY).

### G. lamblia trophozoite in vitro bioluminescence optimization

The optimum concentration of luminescent substrate, D-Luciferin (GoldBio, USA), required for cell identification was determined as earlier described.^10^ AkaLumine (a D-Luciferin analog that is not recognized by the CBG99 enzyme) was used as a control.^15, 16^ Plates were incubated for 5, 10, 15, 30, 60, 90, 120, 150, and 180 minutes at room temperature. Parasite numbers were correlated with bioluminescence intensity by plating a two-fold serial dilution of cells.^10^ Plates were read with an EnVision Multilabel Plate Reader (Perkin Elmer, USA) after incubation at room temperature. Assays were repeated on different days at least twice.

### G. lamblia luciferase expression stability

To evaluate the stability of *G. lamblia* WBC6:pGDHL∷PpyRE9h and *G. lamblia* WBC6:pGDHL∷CBG99 in the absence of puromycin selection, the strains were diluted (1:130 dilutions) twice a week with or without puromycin for 4 weeks. The luciferase activity proportional to cell concentrations was measured as described above.

### G. lamblia cell assay development

Assays were performed in clear, flat bottom 96- or 384-well plates (Corning Inc., Corning, NY) in TYDK media.^10^ Compound screens used one concentration between 20 μM and 1 μM, and dose response assays employed three-fold serial dilutions starting at concentrations up to 60 μM, depending on the particular compound’s potency. All experimental wells were repeated in duplicate or quadruplicate within the same plate. Compounds were incubated with 250,000 *G. lamblia* trophozoites/mL at a final DMSO concentration of ≤0.33%. Negative (DMSO) and positive (Metronidazole) controls were included in each assay. The plates were placed in BD GasPak EZ anaerobe pouches (Becton Dickinson, San Jose, California) and incubated at 37°C for 44 hours. Growth was evaluated under a microscope before vinyl stickers were added to the bottom to prepare for luminescence reading. Ten µL of 2.5 mg/mL D-Luciferin was added to each well and the reaction was incubated at room temperature for 5 minutes on a shaker protected from light. Luminescence was read with an EnVision Plate Reader.^10^ Response curves and EC_50_ values (the concentration at which cell growth is inhibited by 50%) were calculated using GraphPad Prism 6 (GraphPad, LaJolla, CA). All assays were repeated on different days.

Using the *G. lamblia* WBC6:pGDHS∷CBG99 strain, a library of MetRS inhibitors was initially screened at a single concentration of 20 μM. *G. lamblia* WBC6:pGDHS∷CBG99 and WBC6:pGDHS∷CBG/Nluc trophozoites were used to confirm hits in a 2 µM screen, where hits were identified as those causing >80% inhibition of growth.

### Giardia murine model development and efficacy of 1717

Eight- to twelve-week old female BALB/c mice weighing approximately 20 g were used for this study. To promote parasite colonization, mice were administered antibiotics (0.25 mg/mL ampicillin and neomycin) in drinking water for the duration of the experiment starting 6 days before infection.^17^ A cocktail containing ampicillin, neomycin and metronidazole at concentrations of 50 mg/kg, 15 mg/kg and 50 mg/kg, respectively, was dosed by oral gavage 3 days before infection. A second antibiotic dose containing 50 mg/kg ampicillin and 15 mg/kg neomycin was given 1 day before infection. Mice were administered 1×10^7^ *G. lamblia* WBC6:pGDHS∷CBG99 trophozoites in PBS by oral gavage. Establishment of infection was confirmed using a noninvasive IVIS imaging method in which mice were given an intraperitoneal injection of D-Luciferin (150 mg/kg), sedated using isoflurane and placed in the IVIS instrument with nose cones administering continuous anesthetic.^17^

A pilot *in vivo* drug treatment analysis with metronidazole at 50 mg/kg once a day (QD) as a positive control and dosing vehicle (3% ethanol and 7% Tween 80 in normal saline) as a negative control was used for validation of the infection model. This was followed by other pilot studies with **1717** at 50 mg/kg once a day (QD) and dosing vehicle controls to further optimize the model. Once the model was functionally optimized, the following round of experiments tested various doses of **1717**: 50 mg/kg twice a day (BID), 25 mg/kg BID, and 50 mg/kg QD. Control groups included treatment with dosing vehicle and treatment with metronidazole at 50 mg/kg BID. Treatments were introduced by oral gavage for 3 days. The course of infection was tracked by whole-animal imaging.

Fecal samples from individual mice were collected before and throughout the treatment timeline. DNA was extracted from the samples using the QIAamp Fast DNA Stool Mini Kit (Qiagen, Germany). Cyst shedding in treated and untreated mice was verified by PCR analysis of the highly conserved *Giardia* GDH gene using primers and conditions previously described.^18^

### G. lamblia cyst in vitro bioluminescence

An encystation protocol was adapted from previous studies ^19, 20^ to demonstrate the signal intensity of *G. lamblia* WBC6:pGDHS∷CBG/Nluc cysts. Confluent trophozoites were incubated with encystation media for 48 hours, then pelleted and resuspended in growth media for 24 hours. Cells were pelleted and resuspended in cold deionized water and stored at 4°C overnight. The amount of luciferase-induced light emission was quantified as described above, with the modification that Nano-Glo® luciferase reagent (Promega, Madison, WI) was the luminescent substrate.

## Results

### Promoter and luciferase gene selection

We previously used the β-tubulin promoter to drive red-shifted firefly luciferase PpyRE9h in *G. lamblia* WBC6 cells as a tool to assay growth inhibition by compounds in the MMV Pathogen Box.^10^ Further analysis showed that the luminescent signal of PpyRE9h diminished after three weeks of continuous incubation with or without antibiotic selection. To improve the expression level of PpyRE9h, the pβTub was replaced by three Giardia constitutive promoters: a 206 bp pOCT,^12^ a 44 bp pGDHS,^11^ and a 165 bp pGDHL.^11^ The resultant *G. lamblia* trophozoites expressing pGDHL∷PpyRE9h and pOCT∷PpyRE9h had two- and three-fold increased luminescence expression levels relative to pβTub∷PpyRE9h, respectively. There was no significant change for pGDHS∷PpyRE9h relative to the pβTub∷PpyRE9h strains. However, the highest luminescence value that any combination of promoter and PpyRE9h achieved was a modest 1200 photons/sec for the *G. lamblia* WBC6:pOCT∷PpyRE9h strain (Figure 1A). We speculated that this would not be bright enough for use in a mouse model of infection, where the luminescent signal must penetrate multiple layers of tissue to visualize infection noninvasively. There also appeared to be a fitness cost for cells with the pOCT-driven luciferase exhibited by a decrease in growth rate. Thus, further work was needed to establish a brighter and more stable luciferase reporter system in *G. lamblia*. Earlier studies had demonstrated that Fluc, Luc2, and CBR had higher sensitivity than PpyRE9h to report promoter activity in other systems.^21, 22^ Hence, these red luciferases were assayed in *Giardia*.

**Figure 1.**
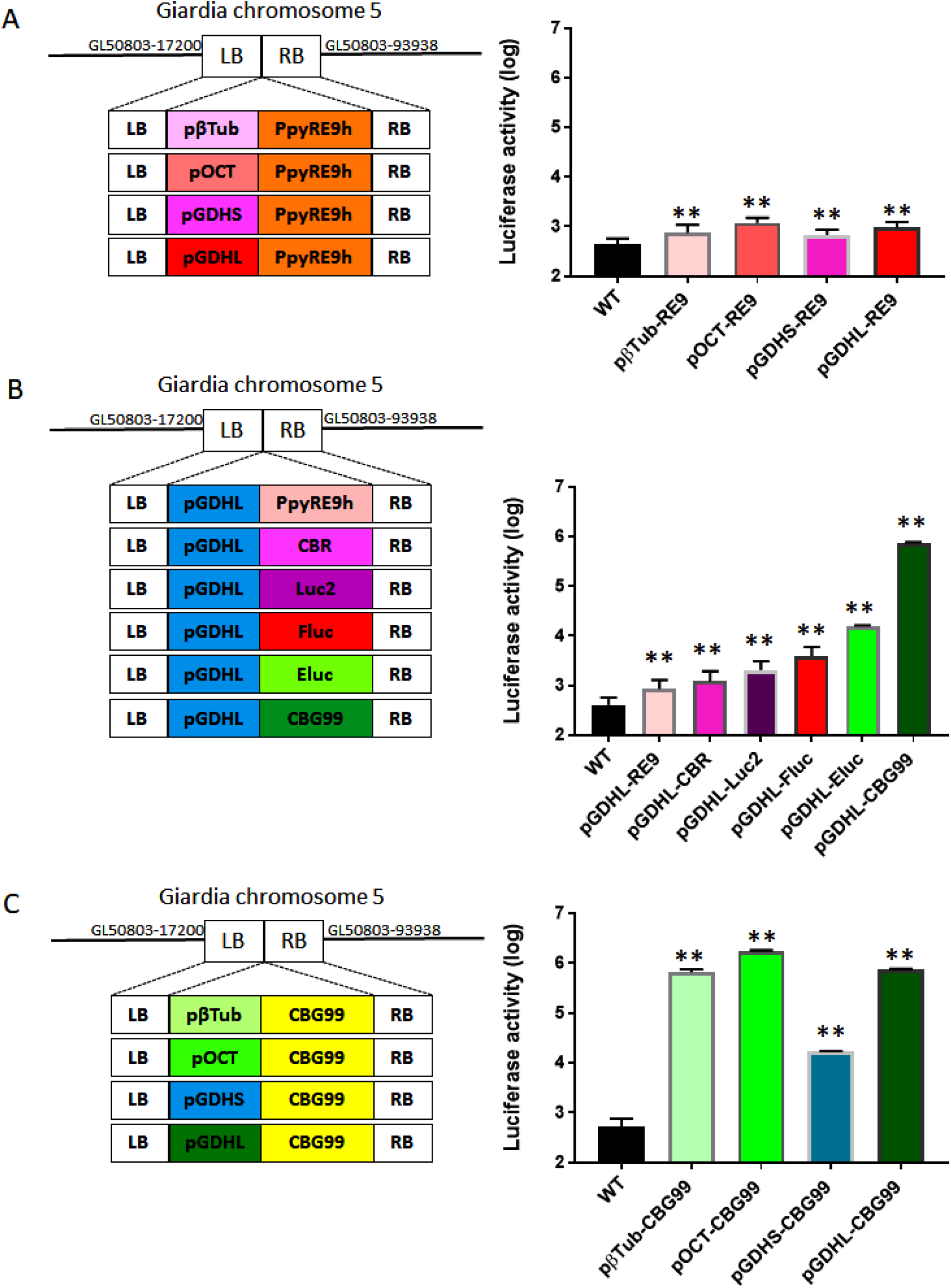
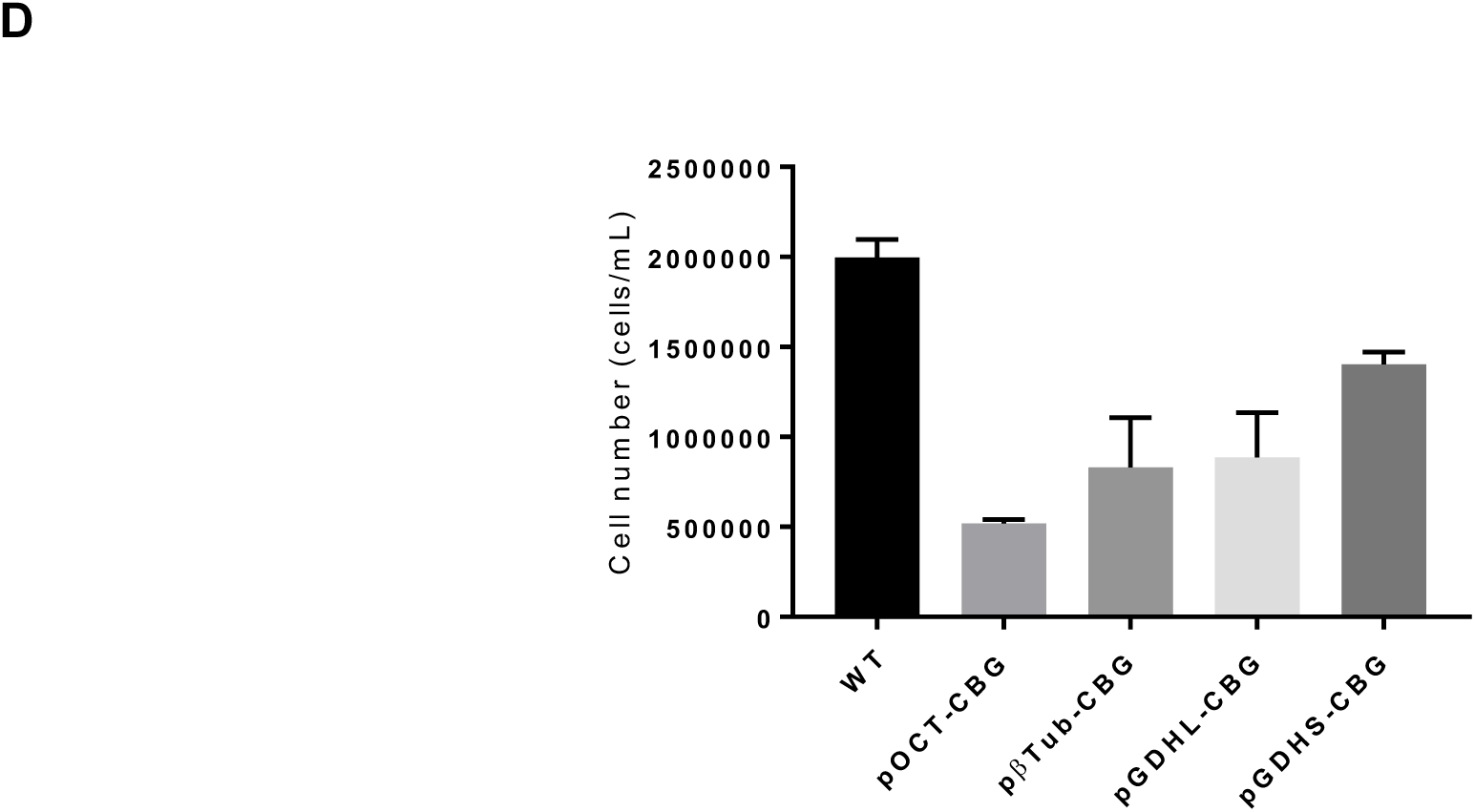
*In vitro* characterization of luciferase sensitivity. (a) Left: Schematic of PpyRE9h driven by four Giardia specific promoters and homologous recombination. LB=left border, RB=right border. Right: *In vitro* comparison of PpyRE9h-dependent photon flux driven by four Giardia specific promoters. (b) Left: Schematic of six different luciferases driven by pGDHL and homologous recombination. Right: *In vitro* quantitative analysis of luciferase-dependent photon flux from six different luciferases driven by pGDHL. (c) Left: Schematic of CBG99 driven by four Giardia specific promoters and homologous recombination. Right: *In vitro* comparison of CBG99-dependent photon flux driven by four Giardia specific promoters. (d) Effect of different promoters on growth rate. At day 1, 150 μL (1×10^5^ cells/mL) of culture was transferred to 13 mL TYDK. Cell density measurement after 3 days incubation at 37°C showed a significant fitness loss for most of the luciferase strains relative to the parent wild type parasite. *G. lamblia* WBC6:pGDHS∷CBG99 showed the lowest fitness disadvantage.

The red luciferase genes PpyRE9h, Fluc, Luc2, and CBR inserted downstream of pGDHL were transfected into *G. lamblia* WBC6 cells. Among the resultant strains, trophozoites with the Fluc gene had the brightest signal intensity, followed by those with the Luc2 gene (Figure 1B). Two green luciferase emitting genes, CBG99 and ELuc, previously shown to exhibit higher levels of bioluminescence than Fluc,^23, 24^ were also assayed for higher signals under the control of pGDHL. In our study, *G. lamblia* WBC6:pGDHL∷CBG99 was 100-fold brighter in signal intensity than *G. lamblia* WBC6:pGDHL∷Eluc. We conclude that *G. lamblia* WBC6:pGDHL∷CBG99 is the brightest luciferin-based reporter system among the red and green luciferases tested (Figure 1B).

A comparative analysis to determine the most suitable promoter for CBG99 in *Giardia* was subsequently performed. The CBG99 gene in the transfection construct was placed under the control of four different *Giardia* promoters: pβTub, pOCT, pGDHS and pGDHL. All the transgenic *G. lamblia* WBC6 strains expressing CBG99 luciferase gave robust bioluminescence signals, with pGDHS∷CBG99 being the lowest even on the log scale (Figure 1C). However, *G. lamblia* strains expressing the CBG99 luciferase driven by pβTub, pOCT and pGDHL promoters have substantial fitness disadvantages in their rate of growth and proliferation relative to the wild type strain. The effect was least pronounced with the *G. lamblia* WBC6:pGDHS∷CBG99 (Figure 1D). Though it had the lowest signal of the promoters tested with CBG99, the signal intensity was still several fold brighter than the one given by pGDHS∷PpyRE9h, and therefore it was more suitable for the drug screening assays and murine model development.

### G. lamblia WBC6:pGDHS∷CBG99 in vitro bioluminescence signal

Optimum luminescence was achieved after 5 minutes of incubation with D-Luciferin at a final concentration of ~83 µg/mL. At this concentration and incubation time, the relative light units signal was high enough to distinguish living cells at 10^3^ cells/mL concentrations. The signal remained stable for up to 30 minutes after the addition of substrate, but the luminescence decreased at longer incubation times. The amount of CBG99-driven bioluminescence activity was directly proportional to the number of viable transfected *G. lamblia* trophozoites per well, as determined by a comparative analysis with the cell count of transfected strains (Figure 2A). The blank media control wells showed no luminescence as compared to the empty background wells and to the sample wells. AkaLumine showed no signal at any time point, as expected. The average Z’ of this assay was 0.6.

**Figure 2:**
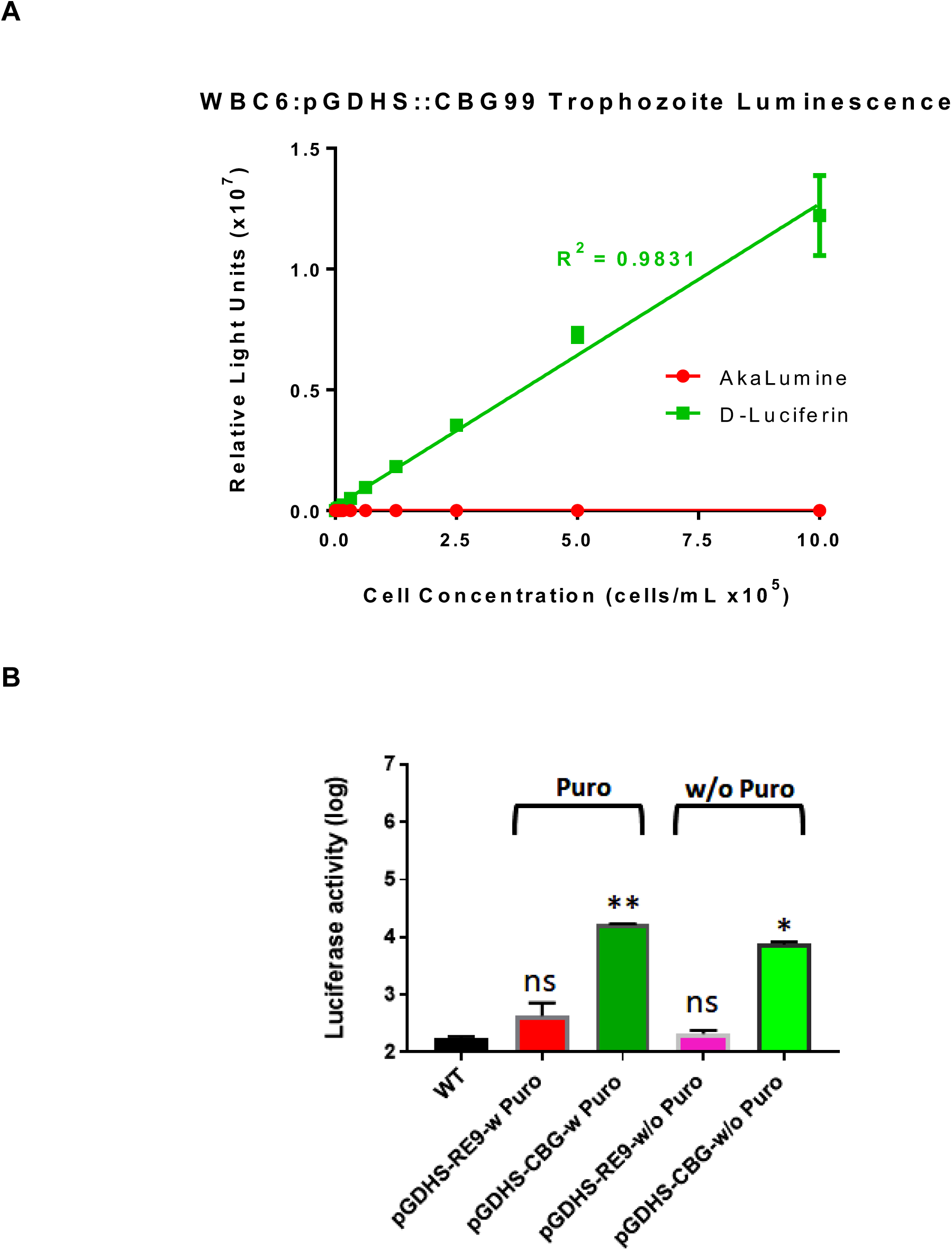
Parasite cell count vs relative light units (RLU). (a) The plot shows the linearity of bioluminescence readouts from the *G. lamblia* WBC6:pGDHS∷CBG99 strain. Readout follows 5 minutes of incubation at room temperature. AkaLumine showed no signal at any time point. (b) *In vitro* comparison of PpyRE9h-dependent and CBG99-dependent photon flux in the presence or absence of selection antibiotic (puromycin) for four weeks. Level of significance is indicated by ns=not significant, * p≤0.001 and ** p≤0.0001.

### G. lamblia luciferase expression stability

In our previous study, the bioluminescent signal of *G. lamblia* WBC6:pβTub∷PpyRE9h was the same as the wild type *G. lamblia* WBC6 after three weeks of incubation, suggesting a loss of the luciferase gene activity. Therefore, we tested for signal stability in the *G. lamblia* WBC6:pGDHS∷CBG99 cells in the absence of the selection antibiotic, puromycin. After four weeks without antibiotic, the bioluminescent signal of *G. lamblia* WBC6:pGDHS∷CBG99 was detected in sufficiently high levels to be called stable expression. The signal was 35 times brighter than that given by *G. lamblia* WBC6:pβTub∷PpyRE9h cells after the same amount of time, which shows no differentiable signal compared to the wild type (Figure 2B). This is especially important to guarantee robust detection of the bioluminescent signal for an *in vivo* experimental model of chronic infection lasting over 3 weeks.

### Screening of MetRS inhibitors including compound *1717*

The initial MetRS inhibitor screen at 20 μM proved to be highly potent against the parasites, with inhibition rates >98% across most compounds. The concentration was subsequently dropped to 2 μM, where most compounds still had potent inhibition (Figure 3A, Table 2). Compounds **BKI1708** and **BKI1770**, which are highly selective inhibitors of apicomplexan calcium dependent protein kinases not found in *G. lamblia*, were included in the screens as negative controls.^25^ The low inhibition rates of **BKI1708** and **BKI1770** at both concentrations demonstrate the specificity of the assay. Percentage inhibition data from the compound screens are presented below (Table 2).

**Table 2:** Percentage inhibition of luciferase *G. lamblia* cells by MetRS inhibitors showed similar inhibition profile for the CBG99 and CBG99/Nluc strains

| Compounds | Chemical structures | Percentage inhibition |  |  |  |
| --- | --- | --- | --- | --- | --- |
|  |  | CBG99 |  | CBG99/Nluc |  |
|  |  | Mean(%) | SEM | Mean (%) | SEM |
| 1331 |  | No inhibition | - | No inhibition | - |
| 1614 |  | No inhibition | - | No inhibition | - |
| 1717 |  | 96 | 0 | 96 | 0 |
| 1849 |  | 4 | 8 | 8 | 7 |
| 1893 |  | 8 | 9 | No inhibition | - |
| 2080 |  | 89 | 1 | 83 | 0 |
| 2093 |  | 97 | 0 | 96 | 0 |
| 2114 |  | 93 | 2 | 97 | 0 |
| 2144 |  | 99 | 0 | 99 | 0 |
| 2207 |  | 95 | 1 | 97 | 1 |
| 2240 |  | 98 | 1 | 96 | 1 |
| 2242 |  | 90 | 1 | 93 | 1 |

**Figure 3A:**
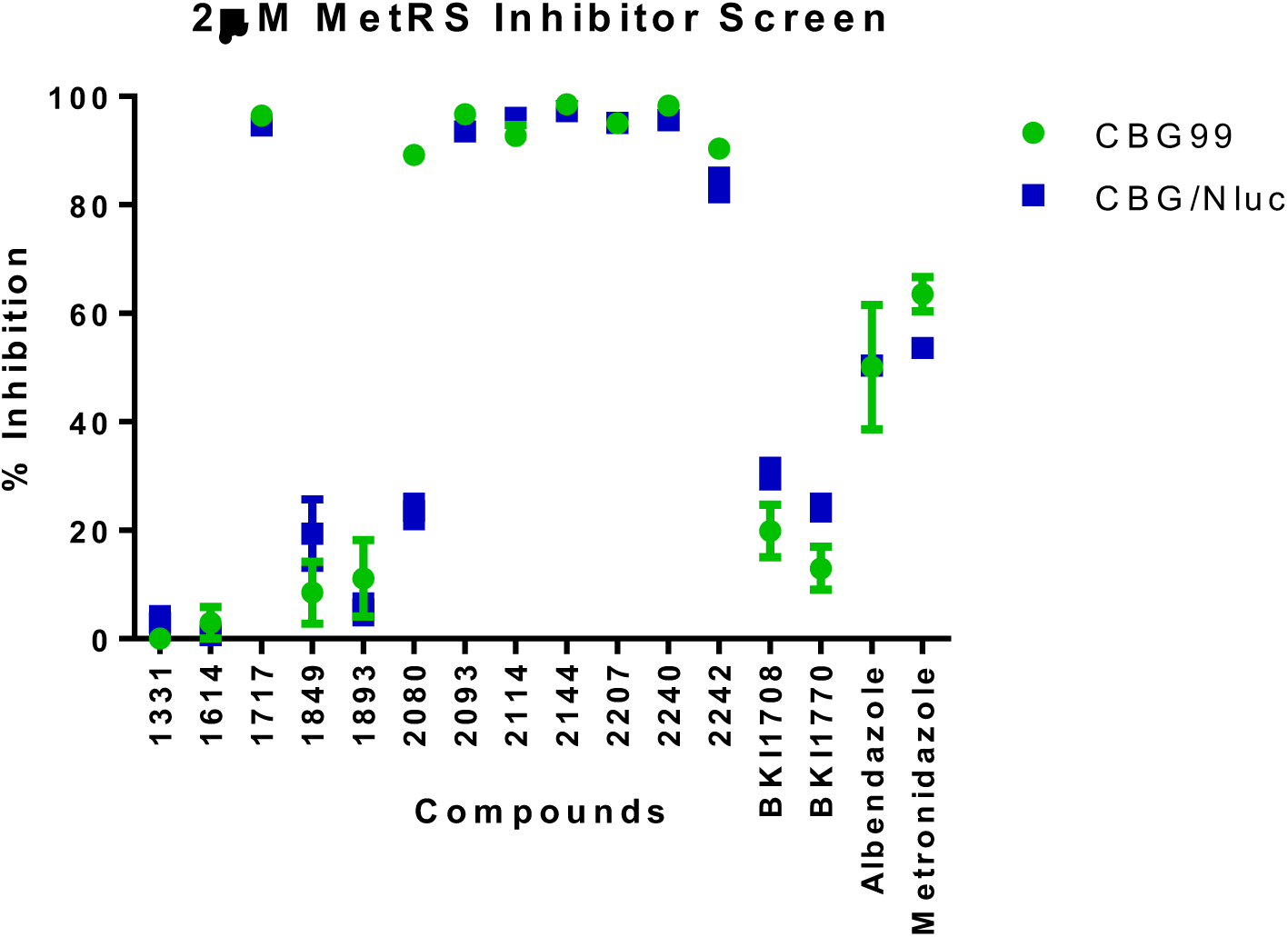
MetRS inhibitor screening of *G. lamblia* luciferase emitting trophozoites of WBC6:pGDHS∷CBG99 versus WBC6:pGDHS∷CBG/Nluc.

**Figure 3B:**
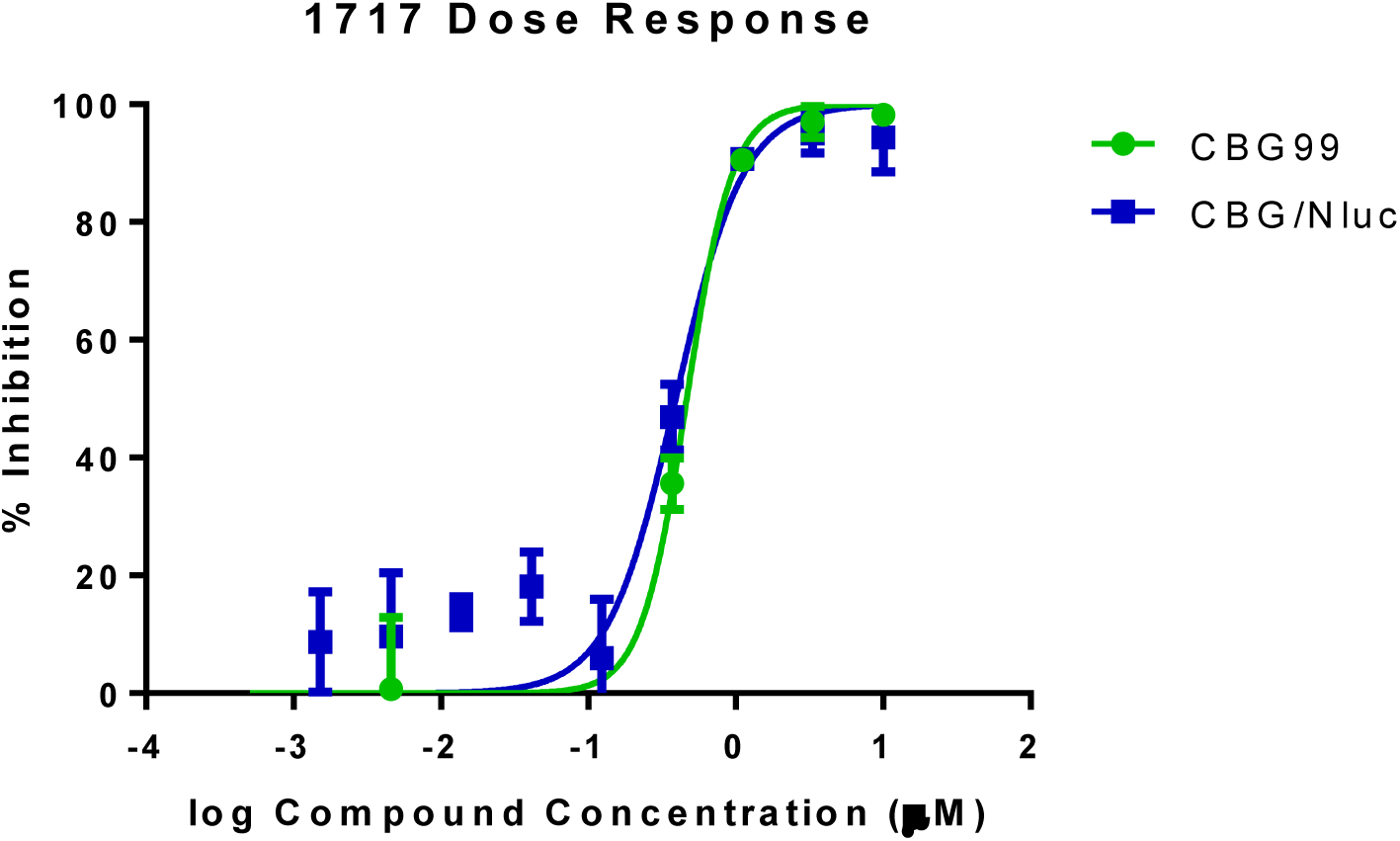
A dose response plot of Compound **1717** inhibition of *G. lamblia* luciferase emitting trophozoites WBC6:pGDHS∷CBG99 versus WBC6:pGDHS∷CBG/Nluc.

The 2 μM screen in 384-well plates with *G. lamblia* WBC6:pGDHS∷CBG99 and *G. lamblia* WBC6:pGDHS∷CBG/Nluc had average Z’ values of 0.7 and 0.6, respectively. The screening data shows that these compounds exhibit a similar inhibition profile against both strains (Figure 3A). Compound **1717** has previously shown potent inhibition on *Giardia* MetRS enzyme (*Gl*MetRS) activity and its EC_**50**_ was determined to be 453 nM.^26^ The EC_**50**_ of **1717** for *G. lamblia* WBC6:pGDHS∷CBG99 and WBC6:pGDHS∷CBG/Nluc was found to be 465 nM and 392 nM, respectively (Figure 3B). The metronidazole EC_**50**_ values of 2.867 µM and 3.114 µM obtained for *G. lamblia* WBC6:pGDHS∷CBG99 and WBC6:pGDHS∷CBG/Nluc strains, respectively, are within the range of previously reported literature values.^10, 26–28^

### Efficacy of metronidazole and 1717 in the Giardia murine model

We describe here the development and evaluation of a mouse infection model as a tool for measuring growth and proliferation of parasites in experimental drug treatment assays. Infection in BALB/c female mice was established within 5 days as determined by IVIS imaging and held for more than two weeks while bacterial antibiotics were maintained in drinking water. The rate of *G. lamblia* WBC6:pGDHS∷CBG99 infection of BALB/c female mice was around 90%, which is consistent with earlier reports.^29^ IVIS image analysis revealed infection signals of up to 100-fold higher radiance values than background luminescence from uninfected mice.

The infection model using the *G. lamblia* WBC6:pGDHS∷CBG99 strain was validated by metronidazole, the current available therapeutic. At 50 mg/kg QD, it delivered a cure after 4 days. In the subsequent round of experiments, the efficacy of compound **1717** was determined. **1717** showed no adverse effects during treatment and all mice were cleared of the infection after 3 days of dosing with 50 mg/kg BID, 25 mg/kg BID, and 50 mg/kg QD. Imaging 24 hr after the final dose and one week after the final dose showed that the drug cleared the infection compared to the vehicle controls (Figure 4). A cure, in this case, is defined as a lack of visible luminescence above background levels in the IVIS images (as confirmed by the plot of the radiance in Figure 4) a week after the last dose and a negative reading from the stool PCR.^30^ Molecular detection analysis with PCR confirmed the absence or presence of *Giardia* cysts expelled in the feces of treated and untreated mice. All procedures involving animals were conducted in adherence to federal regulations and University of Washington’s Institutional Animal Care and Use Committee (IACUC) guidelines.

**Figure 4:**
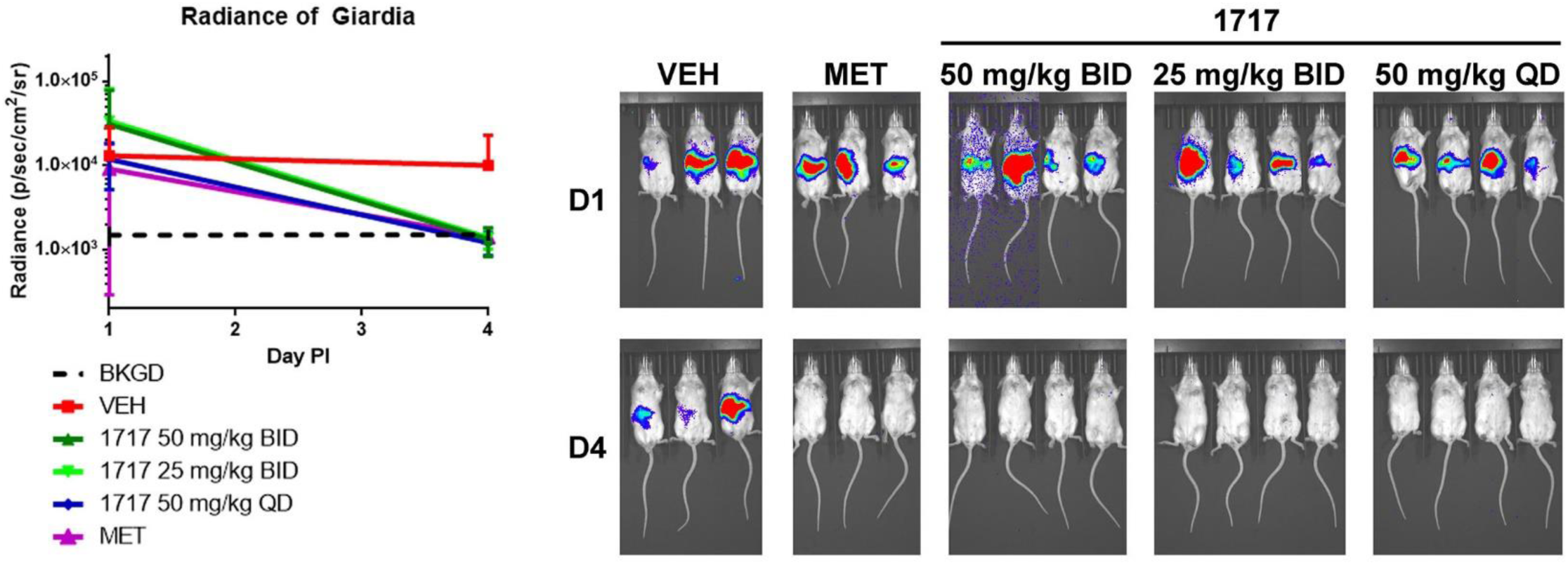
Radiance plot and *Giardia* mouse infections before and after treatment. These are noninvasive imaging of trophozoite growth in mice, 5 days post-infection (D1) with the *G. lamblia* WBC6:pGDHS∷CBG99 strain before the start of treatment. Mice were treated with 50 mg/Kg Metronidazole BID (MET), compound **1717** dosed at 25 mg/kg or 50 mg/kg BID and 50 mg/kg QD. Treatment duration was for 3 days. Mice Panel VEH were dosed with the dosing vehicle (3% ethanol, 7% Tween 80 in normal saline) as a control. Imaging on day 8 post infection (D4) showed that **1717** and metronidazole cleared the infection relative to the untreated controls (VEH). The plot of the intensity/radiance data before therapy against days after start of treatment showed a significant drop relative to untreated controls. * Note that images for some groups of mice on D1 have been cut together. This is due in part to differential intensity of the luminescence signal obtained from some mice hence differences in the length of time needed to acquire images that confirmed established infection. All images were taken during the same session and were scaled together.

### G. lamblia WBC6:pGDHS∷CBG99 versus WBC6:pGDHS∷CBG/Nluc

To further confirm infection and clearance during *in vivo* assays, we attempted to measure luminescence in *G. lamblia* WBC6:pGDHS∷CBG99 cysts from fecal samples. There was no detectable luminescence even in our untreated infected controls. This is likely because the cyst wall, which protects cysts from rupturing in water, prevents the uptake of D-Luciferin. To improve upon this reporter strain, we endogenously tagged the CWP1 gene of the *G. lamblia* WBC6:pGDHS∷CBG99 strain with Nluc. A plot of bioluminescence outputs from cysts of *G. lamblia* WBC6:pGDHS∷CBG99 versus WBC6:pGDHS∷CBG/Nluc using NanoGlo is presented in Figure 5A. The growth and proliferation rate of the 2 strains was experimentally compared and shown to be similar (Figure 5B).

**Figure 5:**
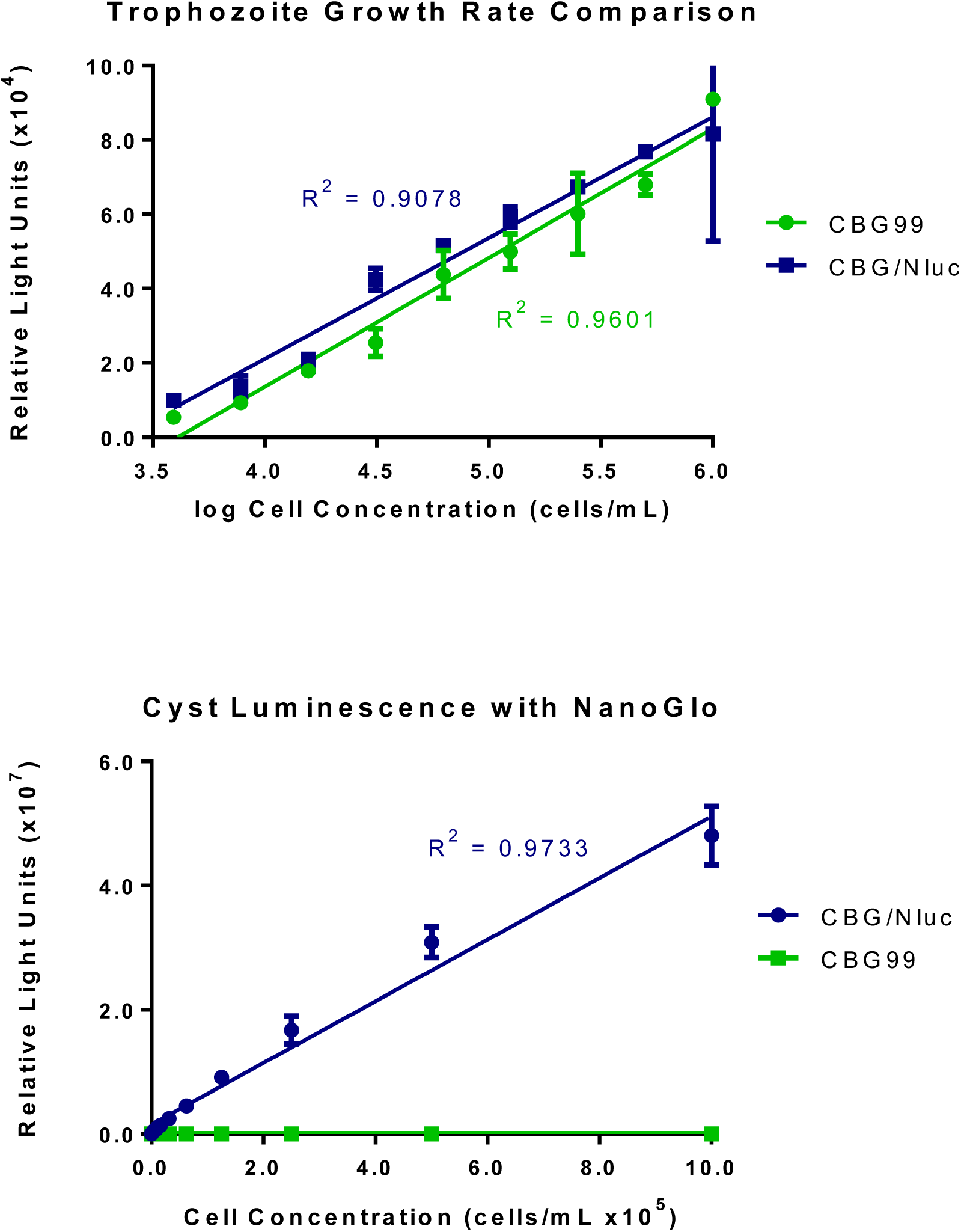
*G. lamblia* WBC6:pGDHS∷CBG99 versus *G. lamblia* WBC6:pGDHS∷CBG/Nluc, (a) A plot comparing the growth rate of *G. lamblia* WBC6:pGDHS∷CBG99 versus *G. lamblia* WBC6:pGDHS∷CBG/Nluc. The same serial concentrations of trophozoites from both strains were incubated at 37°C in *Giardia* growth media. Cell count versus luminescence was plotted after 44 hours. (b) Plots of bioluminescence readouts from dual luciferase *G. lamblia* WBC6:pGHDS∷CBG/Nluc cysts showed a linear relationship to cell counts relative to no signal from the WBC6:pGDHS∷CBG99 strain.

## Discussion and Conclusion

Diarrheal syndromes are among the leading causes of morbidity and mortality by infectious diseases. *G. lamblia*, a significant contributor to the overall global diarrheal burden,^31^ poses a public health challenge due to diminishing treatment options. The luminescent *G. lamblia* strains described here could be an important tool in the search for new giardiasis treatments. *G. lamblia* WBC6:pGDHS∷CBG99 and WBC6:pGDHS∷CBG/Nluc were developed by a process of iterative refinement that led to the selection of the most suitable luciferase and promoter combination. The bioluminescence signal remained stable for weeks *in vitro* and *in vivo* without concurrent administration of puromycin to maintain selective pressure. We earlier determined that Nluc is approximately 17 times brighter than CBG99 when driven by the same promoter (not shown). Since mice shed relatively few cysts during infection, a more sensitive luciferase is required to validate infection or clearance in experimental drug studies, which makes Nluc a good choice. CBG99 is a better cell viability reporter as it requires the presence of ATP for luminescence and therefore will not produce light in dead cells. That Nluc does not require ATP for luminescence makes it a viable reporter for the cyst stage when cells are relatively dormant. Hence, Nluc was not considered as an alternative to CBG99 but rather as a supplement. Additionally, fusing Nluc to CWP1, which is secreted to form the cyst wall, eliminates the need for luminescent substrate to travel through the cyst wall barrier in order to reach the luciferase. This modification will provide the ability to follow total parasite load in animals as well as easily quantify cyst production.

We demonstrated the usefulness of the luciferase reporter system to evaluate treatment with compound **1717** in a mouse model of giardiasis using the noninvasive IVIS Spectrum optical imaging system. Compound **1717**, a fluoro-imidazopyridine, is one of a new class of inhibitors that stop protein synthesis by targeting parasitic MetRS as previously described.^26^ These inhibitors are lethal to *G. lamblia* parasites but nontoxic to mammalian cells in cell based assays.^26^ Chemical synthesis, pharmacokinetics, cytotoxicity, and inhibitory activity of compound **1717** on the wild type *G. lamblia* WBC6 strain and the *Gl*MetRS enzyme were previously described.^26, 32, 33^ Analysis of **1717**’s pharmacokinetic profile showed that a single 50 mg/kg oral dose in mice would have sufficient gut and systemic levels to be effective for treatment of mouse giardiasis.^33^ The compound is about 8 times more potent than metronidazole, which has an EC_50_ of 5 µM.^26^ It has solubilities of 52 μM at pH 7.4, 96 μM at pH 6.5 and 100 μM at pH 2. Earlier experiments suggest that MetRS inhibitor **1717** has a “cidal” anti-*Giardia* activity.^26^ All of these factors lead to the use of **1717** in the present study as a potential therapeutic. Our results further support **1717**, as its efficacy in clearing *Giardia* infections in mice has been demonstrated. Compound **1717**, developed as a trypanosomal agent, was used as a proof of principle molecule to demonstrate that inhibitors of *Gl*MetRS could be a starting point for the development of new anti-giardiasis therapeutics. Further development of this series of inhibitors could facilitate innovative therapeutic options for Giardiasis whose etiologic agent, *G. lamblia*, is rapidly evolving past currently available therapies. Since there is no overlap in mechanism of action with any currently available drugs, cross-resistance is unlikely.

## Acknowledgement

The authors would like to thank Dr. Kelly M. Hennessey, Matthew A. Hulverson, Nora Molasky, Wesley C. Van Voorhis, and Ryan Choi for helpful discussions.

## Funding

This study was supported by grants from National Institute of Allergy and Infectious Diseases and National Institute of Child Health and Human Development under award numbers R01AI110708, R01AI097177, R21AI119715, R21AI127493, R21AI123690 and R21AI140881.

## Transparency declarations

None to declare.

